# Switching of OAS1 splicing isoforms mitigates SARS-CoV-2 infection

**DOI:** 10.1101/2021.08.23.457314

**Authors:** Kei Iida, Masahiko Ajiro, Yukiko Muramoto, Toru Takenaga, Masatsugu Denawa, Ryo Kurosawa, Takeshi Noda, Masatoshi Hagiwara

## Abstract

**Background:** The rapidly accumulating disease susceptibility information collected from coronavirus disease (COVID-19) patient genomes must be urgently utilized to develop therapeutic interventions for SARS-CoV-2 infection. Chromosome 12q24.13, which encodes the 2’-5’-oligoadenylate synthetase (OAS) family of proteins that sense viral genomic RNAs and trigger an antiviral response, is identified as one of the genomic regions that contains SNPs associated with COVID-19 severity. A high-risk SNP identified at the splice acceptor site of *OAS1* exon 6 is known to change the proportions of the various splicing isoforms and the activity of the enzyme.

**Methods:** We employed *in-silico* motif search and RNA pull-down assay to define a factor responsible for the *OAS1* splicing. Next, we rationally selected a candidate for slicing modulator to modulate this splicing.

**Results:** We found that inhibition of CDC-like kinase with a small chemical compound induces switching of *OAS1* splice isoforms in human lung cells. In this condition, increased resistance to SARS-CoV-2 infection, enhanced RNA degradation, and transcriptional activation of interferon β1, were also observed.

**Conclusions:** The results indicate the possibility of using chemical splicing modifiers aided by genome-based precision medicine to boost the innate immune response against SARS-CoV-2 infection.

## Introduction

As of July 2021, more than 180 million people have been infected with SARS-CoV-2 and more than 4.0 million people have died of COVID-19 (https://covid19.who.int). Despite extensive efforts towards prophylactic vaccination against SARS-CoV-2, the incidence of COVID-19 continues to increase, with rapid emergence of mutant strains (https://www.who.int/). To understand SARS-CoV-2 pathogenesis and develop novel therapeutic approaches, it is essential to analyze genetic factors responsible for susceptibility and severity. A previous genome-wide association study (GWAS) in over 2,000 COVID-19 patients uncovered several susceptibility-associated genes, including *LZTFL1, CCHCR1, OAS1, OAS2, OAS3, DPP9, TYK2*, and *IFNAR2* (Pairo-Castineira et al., 2021). The COVID-19 host genetics initiative also gathered genetic information for more than 30,000 COVID-19 patients, including over 5,000 patients with very severe respiratory symptoms, and showed the similar results (The COVID-19 Host Genetics Initiative, 2021). It revealed 131 SNPs associated with severe respiratory conditions (p < 5 × 10^−8^) in the gene cluster of *OAS1, OAS2*, and *OAS3* (Fig. 1a). They consist of 5 SNPs within the *OAS1* locus, 72 SNPs within intergenic regions across *OAS1, OAS2*, and *OAS3*, 46 SNPs within the *OAS3* locus, and 8 SNPs within the *OAS2* locus. Importance of the *OAS* family of genes was further indicated in a transcriptome study in an *in vitro* model of SARS-CoV-2 infection (Blanco-Melo et al., 2020), in which normal human bronchial epithelial (NHBE) cells exhibited an increased expression of *OAS1, OAS2*, and *OAS3* (2.3, 2.1, and 2.2-fold increase, respectively). Of the aforementioned SNPs, this genomic region includes SNPs associated with hospitalized COVID-19 patients (32 SNPs) and all COVID-19 patients (125 SNPs) with low p-values (p < 5 × 10^−8^) (Fig. 1b, 1c), which are concentrated around the terminal exon of *OAS1* and intron 2 of *OAS3*. Amongst them, the SNP with the lowest p-value was the G>A SNP rs10774671, located at the last base of intron 5 (Table 1), and its association with infectious viruses, including SARS coronavirus and Dengue virus, has been reported (Hamano et al., 2005; He et al., 2006; Lin et al., 2009).

**Table 1.** SNPs on *OAS1* locus associated with very severe respiratory confirmed COVID-19 (p < 5 × 10^−8^).

| Chr | Pos | ID | p-value* | log<br>(Odds<br>Ratio)<br>* | Ref | Alt | Location to OAS1 transcript<br>variant 1 (p46) |
| --- | --- | --- | --- | --- | --- | --- | --- |
| 12 | 112,912,991 | rs2057778 | $6.86 \times 10^{-10}$ | 0.17 | G | T | Intron 3 (1756th base of<br>5273nt.) |
| 12 | 112,914,354 | rs4767023 | $1.70 \times 10^{-10}$ | 0.17 | T | C | Intron 3 (3119th base of<br>5273nt.) |
| 12 | 112,919,388 | rs10774671 | $1.10 \times 10^{-12}$ | 0.18 | G | A | Intron 5 (1688th base of<br>1688nt.) |
| 12 | 112,919,404 | rs1131476 | $4.54 \times 10^{-12}$ | 0.18 | G | A | Exon 6 (16th base of 515nt.,<br>CDS, Ala>Thr) |
| 12 | 112,919,637 | rs2660 | $2.68 \times 10^{-12}$ | 0.19 | G | A | Exon 6 (249th base of 515nt.,<br>3' UTR) |
| * From GWAS analysis for very severe respiratory confirmed COVID-19 |  |  |  |  |  |  |  |

**Figure 1.**
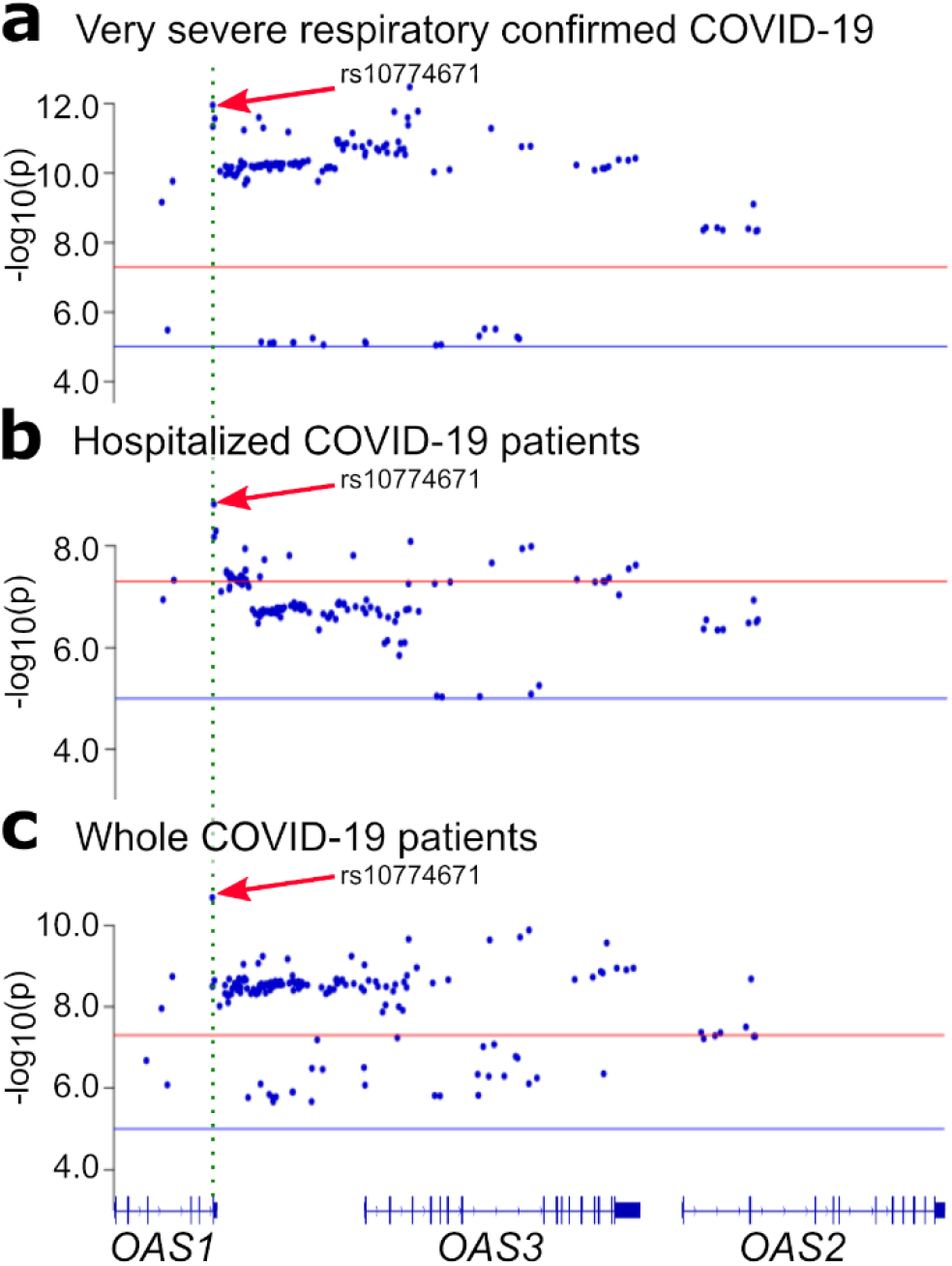
*OAS1-3* loci have COVID-19 associated SNPs and respond to SARS-CoV-2. Positions and GWAS p-values for SNPs on *OAS1, OAS3*, and *OAS2* gene loci. GWAS results for confirmed COVID-19 cases with very severe respiratory conditions (a), hospitalized COVID-19 patients (b), and all COVID-19 patients (c), reported by COVID-19 Host Genome Initiative groups, are shown (Reported by COVID-19 Host Genome Initiative groups). Red and blue horizontal lines show the p-value positions of 5E-8 and 1E-5, respectively. The position of rs10774671 SNP is shown with a green dotted line and red arrows.

## Materials and Methods

### Analysis of GWAS data

GWAS results were obtained from the COVID-19 Host Genetics Initiative data. The following terms were used: “A2_ALL_leave_23andme” for GWAS of confirmed COVID patients with very severe respiratory conditions vs. population, “B2_ALL_leave_23andme” for GWAS of hospitalized COVID patients vs. population, and “C2_ALL_leave_23andme” for GWAS of all COVID patients vs. population. Data version round 5 was used for all the data obtained in this study (January 18, 2021). The sample size of 5582, 12888, and 36590 individuals were chosen for evaluation of GWAS datasets of severe respiratory symptoms, hospitalization, and COVID-19 susceptibility, respectively, obtained from COVID-19 host genome initiative database (The COVID-19 Host Genetics Initiative, 2021).This sample number was determined based on number of applicable data without exclusion, and a sample-size estimation was not conducted prior to the analysis.

### Cell lines

Daudi cells, derived from Burkitt’s lymphoma, were obtained from the cell bank of the National Institutes of Biomedical Innovation, Health and Nutrition (NIBIOHN, Osaka, Japan) and maintained in RPMI 1640 medium (Nacalai Tesque, Kyoto, Japan) supplemented with 20% fetal bovine serum, 100 U/mL penicillin, and 100 µg/mL streptomycin. Calu-3 cells, derived from lung adenocarcinoma, were obtained from American Type Culture Collection (ATCC, Manassas, VA, USA), and cultured in Dulbecco’s modified Eagle’s medium (DMEM) (Nacalai Tesque) supplemented with 10 % fetal bovine serum, 100 U/mL penicillin, and 100 µg/mL streptomycin. All cells were maintained in an incubator at 37 °C with 5 % CO_2_, and mycoplasma was confirmed negative in routine polymerase chain reaction tests. VeroE6/TMPRSS2 cells were obtained from JCRB Cell Bank, and cultured in Dulbecco’s modified Eagle’s medium (DMEM) (Sigma-Aldrich) supplemented with 10 % fetal bovine serum, 100 U/mL penicillin, and 100 µg/mL streptomycin. The cells were maintained in an incubator at 37 °C with 5 % CO2.

### RNA pull-down assay and western blot

Calu-3 whole cell lysates were prepared using a lysis buffer containing 10 mM Tris-HCl (pH 7.4), 150 mM NaCl, 1 mM ethylenediaminetetraacetic acid, 1% Triton X-100, 0.1% sodium dodecyl sulphate, 0.25% sodium deoxycholate, and 10% glycerol with protease inhibitors (Nacalai Tesque) and phosphatase inhibitors (Sigma-Aldrich, Munich, Germany), followed by sonication, treatment with DNase I (Promega, Madison, WI, USA) at 37 °C for 5 min, and centrifugation (24,000 *x g* at 4 °C for 15 min). The supernatant was used as the soluble fraction for the RNA-pull down assay. For the RNA pull-down assay, 5′-biotin, 3′-dTdT-attached RNA, designed for the sequence adjacent to the *OAS1* exon 5 splice donor (5′-CUGCUGGUGAGACCUCCUGCUUCC-3′ (oAM685), was incubated with NeutrAvidin beads (Thermo Fisher Scientific, Waltham, MA, USA) for 2 h at 4 °C (no bait RNA was used for the negative control), followed by wash with 1X TBS thrice. RNA-bound NeutrAvidin beads were then incubated with the Calu-3 cell lysate in the presence of 1% DMSO or 10 μM CaNDY for 16 h with rotation at 4 °C. This was followed by washing thrice with tris-buffered saline and elution with Laemmli buffer. Three technical replicates for RNA pull-down assay were independently conducted, three times from a stocked cell lysate. Eluted proteins were then analyzed by western blotting with anti-U1-70k mouse monoclonal antibody (9C4.1) (05-1588, Merk Millipore, Burlington, MA, USA) at a dilution of 1:500 for the detection of U1-70k, anti-SR protein (1H4G7) mouse monoclonal antibody (33-9400, Thermo Fisher Scientific, Waltham, MA, USA) at a dilution of 1:200 for phosphorylated SRSF6, and anti-β-actin (ACTB) mouse monoclonal antibody (Ac-15) (sc-69879, Santa Cruz Biotechnology, Dallas, TX, USA) at a dilution of 1:4,000 for ACTB. Chemiluminescent signals were detected using a ChemiDoc MP Imaging System (Bio-Rad, Hercules, CA, USA).

### Transcriptome analysis for CaNDY-treated Calu-3 cells

RNAs were extracted using RNeasy Mini kit (QIAGEN, Hilden, Germany) from Calu-3 cells, treated with 10 µM CaNDY or 0.1% DMSO for 18 h, and applied for RNA-Seq analysis. Three technical replicates for RNA-Seq, where each experiment was independently conducted three times from a stocked total RNA. RNA-seq reads were mapped to the human genome sequences (GRCh38) using STAR (ver. 2.7.1a, https://github.com/alexdobin/STAR) with ENCODE options, using the Ensembl genome annotation (ver. 102). Raw reads were counted with bam files, and TPM values were calculated using RSEM v1.2.31 (https://github.com/deweylab/RSEM). Differentially expressed genes were identified using the method described above. Differential alternative splicing (DAS) events were analyzed with the method previously described (Sakuma et al., 2015) using the rMATS program (http://rnaseq-mats.sourceforge.net/rmats4.1.1/). DAS was defined by the following criteria: FDR < 0.01, read counts ≥ 15, and delta Percent Spliced-In (PSI) ≥ 0.05. We compared the DAS events with the gene annotation information, then classified the event types into events on the exons constituting the productive mRNAs, events on additional exons/regions of the productive mRNAs, and others. For characterizing gene set and transcriptome profiles, we used the Metascape website (https://metascape.org/) and Gene Set Enrichment Analysis (GSEA, https://www.gsea-msigdb.org/gsea/). The same samples were also used for RT-PCR.

### SARS-CoV-2 infection

Calu-3 cells were pre-treated with 10 μM CaNDY or 0.1% DMSO for 24 h, infected with SARS-CoV-2 (SARS-CoV-2/Hu/DP/Kng/19-027) at a multiplicity of infection of 0.01, and maintained in the presence of 10 μM CaNDY or 0.1% DMSO. RNA samples were collected from the cells 24 h post-infection (pi), and the virus titers were determined by the 50% tissue culture infectious dose (TCID_50_) using VeroE6/TMPRSS2 cells at 48 h pi. Titer assay was conducted in six biological replicates for independent cell cultures.

### Analysis of RNA degradation

RNA samples were diluted to 200 ng/ μL. The quality of the diluted RNA samples was evaluated using the Agilent RNA 6000 Nano Kit and the Agilent 2100 Bioanalyzer. For the infected cells, three biological replicates with independent cell cultures were prepared for DMSO and CaNDY treatments, respectively, with the method described above. Total RNA from transcriptome analysis represented uninfected control for this study. The gel-like image of the 2100 Bioanalyzer result was visualized using the 2100 Expert Software (ver. B.02.11, Agilent Technologies) in pseudo colors with default settings. For detailed analyses, the migration time and fluorescence unit data were extracted in CSV format as aligned with the sample data. Next, the fluorescence unit values were normalized to an RNA concentration of 300 ng/μL. We selected the data located between 43.3 s and 48.0 s of the migration time as cleaved RNA products and those between 49.9 s and 51.4 s as the 28S rRNA. The sums of the fluorescence unit values were used for further analysis. For drawing the bar plot, the values were scaled to the mean values of DMSO samples.

### RT-PCR

Total RNA extracted from cultured cells was used for reverse transcription using the PrimeScript RTase (Takara Bio, Shiga, Japan) with random hexamers, and the products were then amplified with ExTaq DNA polymerase (Takara Bio) with target-specific primer sets. Primers used in RT-PCR are listed in Table S1. Detection of RT-PCR products was conducted using the ChemiDoc MP Imaging System (Bio-Rad), with subsequent analysis by Image Lab software (Bio-Rad). RT-PCR was conducted in three technical replicates for a total RNA sample.

### Modelling of the viral infection

The statistical simulation of SARS-CoV-2 infections was performed with the SIR (Susceptible, Infectious, or Recovered) model (Harko et al., 2014) using the “deSolve” package (Soetaert et al., 2010) in the R environment. We assigned the initial number of susceptible persons as 9999 and infectious person as 1. Beta, a parameter for infection rate per day, per person, was set as 0.5 or 0.28 (i.e., 0.5 × 0.56), and Gamma, a parameter for the removal or the recovery rate per day, per person, was set as 0.1, according to a report on the spread of SARS-CoV-2 in April 2020, in the USA (Adam, 2020).

### Data Availability

The datasets generated during and/or analyzed during the current study are available from the corresponding author on reasonable request. The original RNA-seq data were deposited at the Gene Expression Omnibus (GEO) of National Center for Biotechnology Information (NCBI) with the accession ID GSE174398.

## Results

### OAS1 splice variant regulation by SRSF6

The rs10774671 A/G SNP is known to control the production of *OAS1* splice variants. The G allele of rs10774671 (G-allele) creates the AG-dinucleotide that is essential for the recognition of the exon 6 splice acceptor site for *p46* variant production. The A allele of rs10774671 (A-allele) leads to the alternative splicing of *OAS1* pre-mRNA to produce *p42, p48, p44a*, and *p44b* variants (Noguchi et al., 2013) (Fig. 2a). OAS1 protein senses the double-stranded RNA structure, including RNA duplication intermediates of SARS-CoV-2 (Sadler and Williams, 2008; Schlee and Hartmann, 2016), to synthesize 2′-5′-oligoadenylates, which in turn trigger the activation of latent ribonuclease L (RNase L) for viral RNA degradation (Sadler and Williams, 2008; Schlee and Hartmann, 2016). Previous studies revealed that the catalytic activity of OAS1 varies with its splice variants; p46, the major isoform of OAS1 produced in the presence of the G-allele, presents optimal activity, while p42, the major isoform produced in the presence of the A-allele, shows poor activity (Carey et al., 2019; Di et al., 2020).

**Figure 2.**
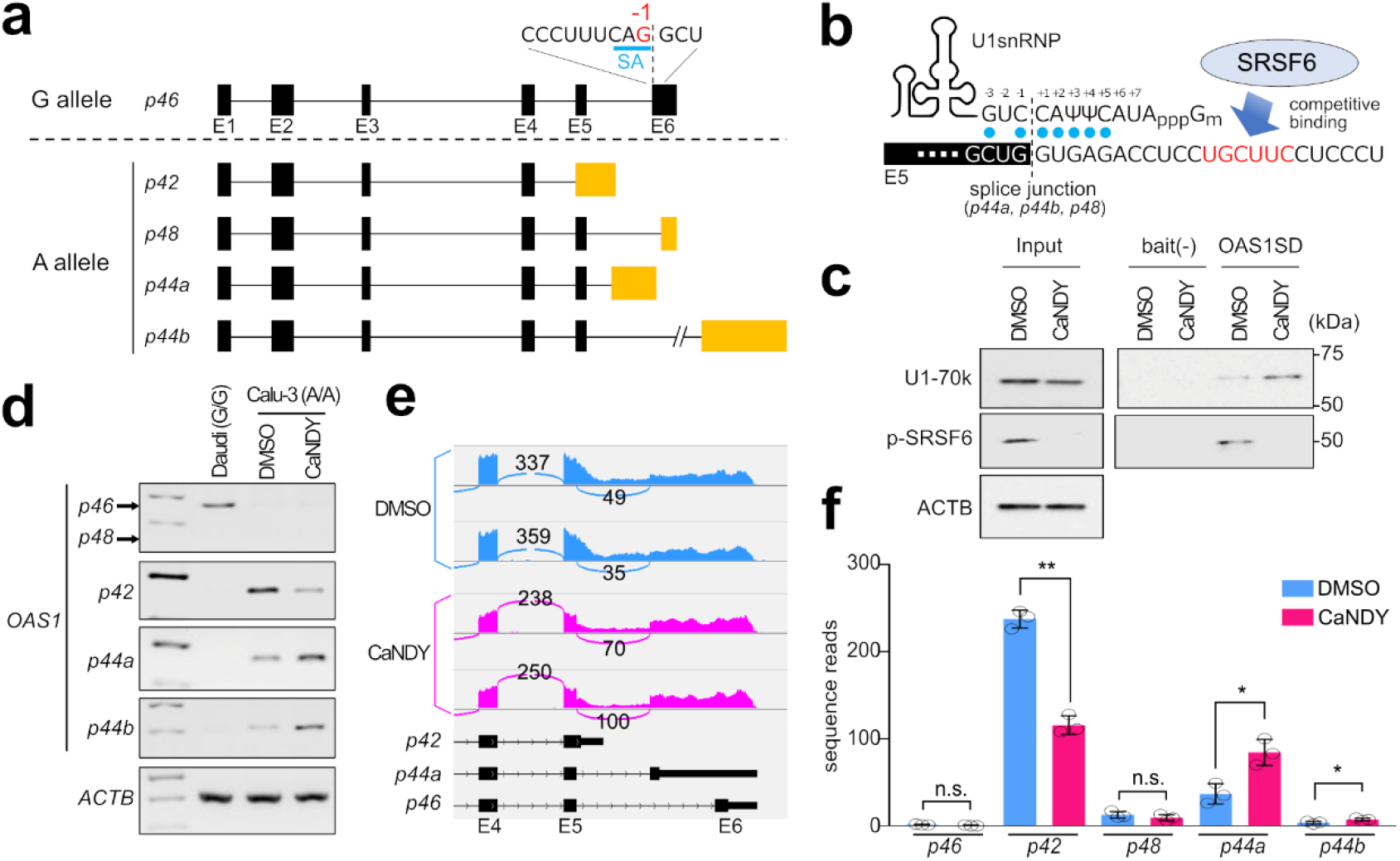
CLK inhibitor CaNDY induces splice-switching in *OAS1* rs10774671 A allele. **a**, Allele-dependent *OAS1* alternative splicing. Pre-mRNA of *OAS1*, with the rs10774671 G-allele at - 1 position of exon 6 splice acceptor (SA), dominantly produces the *p46* variant, whereas the A-allele leads to alternative splicing to produce the *p42, p48, p44a*, and *p44b* variants with differences in the last exon (yellow boxes). **b**, The SRSF6 binding motif (red letters) present in close vicinity of the U1 binding site of *OAS1* donor site for *p44a*/*p44b*/*p48* splicing. Blue dots indicate U1snRNA pairing. **c**, Western blot of U1-70k and phosphorylated SRSF6 (p-SRSF6) for RNA pull-down products of *OAS1* exon 5 splice donor (SD) sequence in Calu-3 cells treated with 0.1% DMSO or 10 µM CaNDY. Input, input samples; bait (-), negative control products without the bait RNA oligo. ACTB served as the loading control. **d**, RT-PCR for *OAS1* alternative splicing profile in Daudi (G/G allele) and Calu-3 (A/A allele) cells treated with 0.1% DMSO or 10 µM CaNDY for 24 h. *ACTB* served as a loading control. **e**, RNA-Seq results indicated by a Sashimi plot for a region covering *p42, p44a*, and *p46*. **f**, Numbers of splice-junction reads for each *OAS1* splice variants. Dots, read number of repeats; error bars, ±SD. n.s., p ≥ 0.05; *, p < 0.05; **, p< 0.01 by Student’s t-test. **Figure 2-source data 1 (Separated File)** Original Western blot files for Figure 2C. **Figure 2-source data 2 (Separated File)** Original gel electrophoresis files for Figure 2D.

In A-allele individuals, disruption of the exon 6 splice acceptor site induces *OAS1* alternative splicing, yielding *p44a, p44b*, and *p48* through activation of acceptor sites downstream of the exon 5, and *p42* as a consequence of the skipped recognition of exon 5 donor site (Fig. 2a). Assuming that the splice-shift from the inactive *p42* to other variants may improve the immune-response that is weakened due to the presence of *OAS1* A-allele, we first looked into the mechanism by which *p42* variant is dominantly produced. In searching for splice motifs surrounding the *OAS1* exon 5 donor sites with the ESEfinder tool (Cartegni et al., 2003), we noticed a binding motif for serine/arginine-rich splicing factor 6 (SRSF6) (5′-UGCUUC-3′) immediately downstream to the exon 5 donor site (Fig. 2b). We then speculated that SRSF6 interaction with this site could prevent U1snRNP from binding to the exon 5 donor site. To test this hypothesis, we applied a pan-CDC-like kinase (CLK) inhibitor, CaNDY, to dissociate SRSF6 from RNA by preventing the kinase activity of CLK to phosphorylate RS domain at the carboxyl-terminal of SRSF6 (Shibata et al., 2020). Following an RNA pull-down assay, we found that the U1snRNP binding to exon 5 donor site was enhanced when Calu-3 cells were treated with CaNDY, as represented by the detection of the U1-70k major subunit in the pull-down products, whereas phosphorylated SRSF6 not detected (*OAS1SD* pull-down products in Fig. 2c). Together, these data indicate that SRSF6 plays a major role in the exon 5 donor site recognition, which is crucial for *OAS1* alternative splicing.

### CaNDY induces splice shift in A-allele

Next, we investigated the consequence of CLK inhibition on alternative splicing of *OAS1* mRNA with A-allele. Calu-3 cells, lung adenocarcinoma cells homozygous for *OAS1* A-allele (Fig. S1a), were treated with 10 µM CaNDY or DMSO and analyzed for *OAS1* splicing profiles, along with Daudi cells, which are lymphoma cells homozygous for the G-allele (Fig. S1b). Consistent with previous observations (Kjær et al., 2014; Noguchi et al., 2013), we found that the *p46* variant was dominantly expressed from *OAS1* G-allele, while the *p42* variant was expressed from the A-allele (Fig. 2d). Intriguingly, CLK inhibition induced splice-switching for *OAS1* with the A-allele, by suppressing *p42* splicing while promoting *p44a* and *p44b* variant production. This observation is consistent with the model suggested by the RNA pull-down assay (Fig. 2c), where the exon 5 donor site is made available for U1snRNP binding upon CLK inhibition (Fig. 2b). However, the *p48* variant expression was under the detection limit in both A- and G-alleles (Fig. 2d). The splicing shift of *OAS1* with A-allele upon CLK inhibition was also evident in RNA-Seq analysis of Calu-3 cells, in which the *p42* variant was suppressed by 51%, and *p44a* and *p44b* were increased by 130% and 100%, respectively; the *p44b* variant was found to be a minor form of *OAS1*, accounting for 1-4% of *OAS1* transcripts (Fig. 2e, 2f). Additionally, through the RNA-Seq analysis of Calu-3 cells with or without CaNDY treatment, we identified differential expression of 98 genes, and splice alterations yielding protein-coding variants for 63 genes; however, none of these events were associated with viral infection except for the *OAS1* splice alterations (Tables S2, S3, and Fig. S2).

### SARS-CoV-2 resistance with splice shift

We examined the alternation of SARS-CoV-2 infection rate in Calu-3 cells, when the splicing isoform was changed from *p42* to *p44a* by manipulating the splicing phenomenon with a small chemical compound. To sufficiently change the balance of the splicing isoforms, cells were pre-treated with CaNDY for 24 h. Viral titer was assayed 2 days after cells were exposed to the virus (Fig. 3a). The virus titer decreased from 5.18 × 10^7^ to 2.90 × 10^7^ (fold change = 0.56) in the cells treated with CaNDY (Fig. 3b, Table S4). There are no reports on the enzyme activity or antiviral activity of the p44a variant; however, based on our results, it can be assumed that the product encoded by *p44a* has stronger enzyme activity/antiviral ability than that encoded by the *p42* form. Furthermore, RNA degradation products were measured to confirm whether this change was indeed due to the switching of *OAS1* splicing isoforms. If enzyme activities of the product encoded by *OAS1* are elevated, elevated activities of RNase L are expected to be observed. Activated RNase L can not only degrade the foreign RNA molecules, but also the endogenous ribosomal RNA within the host cells (Lin et al., 2009). Thus, degradation of rRNA was investigated as a measure of RNase L activity. We confirmed the appearance of a band that is expected to be a degradation product of rRNA in cells infected with SARS-CoV-2 (Fig. 3c). We also confirmed that the concentration of decomposition products, sized 2,000–3,000 nucleotides, increased slightly (fold change = 1.35), with a decrease in the molecular weight and concentration of the 28S rRNA peak (fold change = 0.56), upon CaNDY treatment (Fig. 3c, Fig. S3). It is known that degraded dsRNA activates the transcription of the interferon genes through the OASL and RIG-I related pathways (Choi et al., 2015; Ibsen et al., 2015). We confirmed that CaNDY treatment increases the transcription of *interferon β1* (*IFNB1*) during viral infection (Fig. 3d, 3e). This result was consistent with the previous reports showing *IFNB1* transactivation during viral infections (Ford and Thanos, 2010; Schwanke et al., 2020).

**Figure 3.**
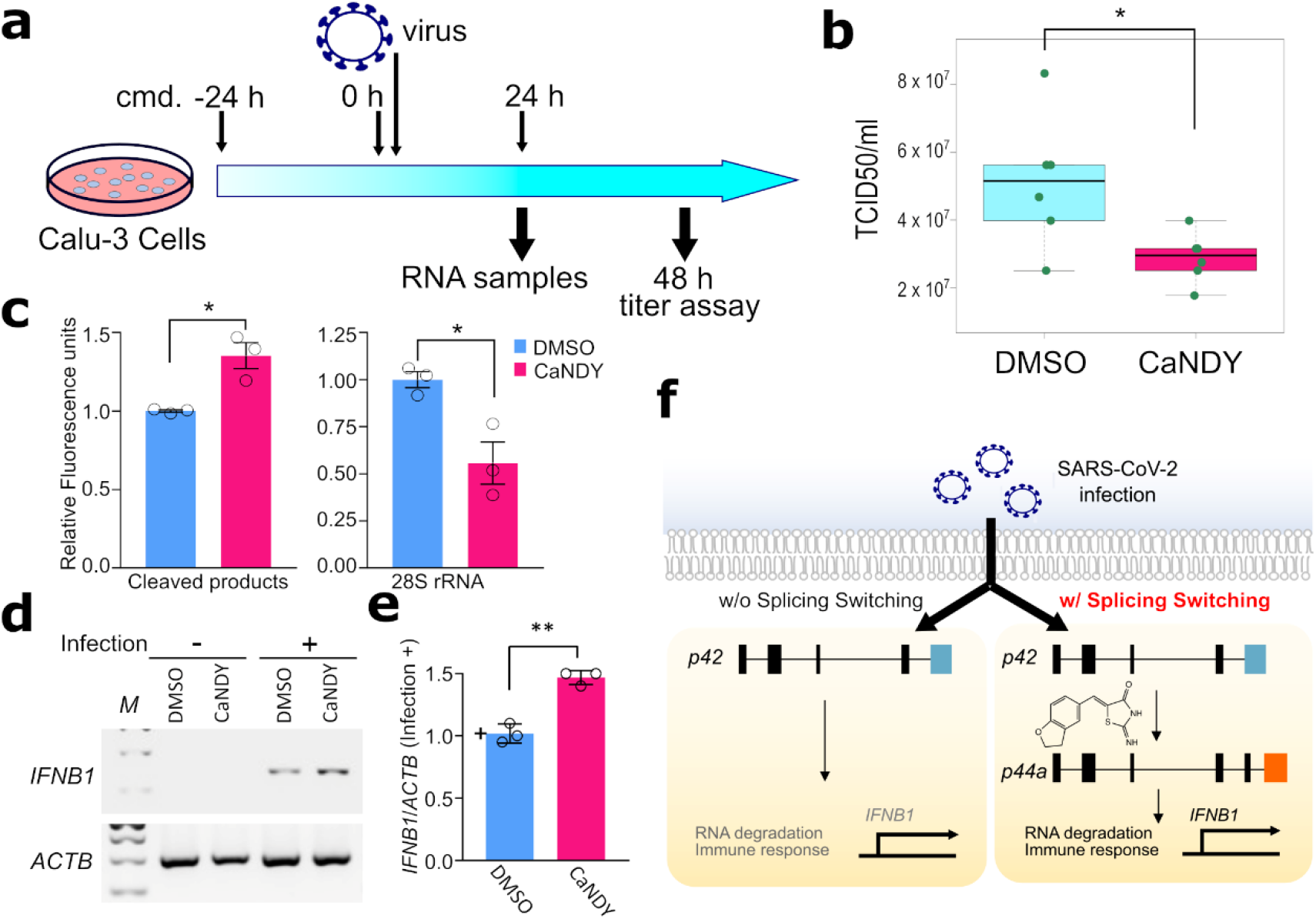
Pre-treatment with CaNDY confers Calu-3 cells with resistance against SARS-CoV-2 infection via activation of the RNase L and interferon pathways. a, A scheme for the pre-treatment of CaNDY, virus titer assay. b, Box plots showing logarithm translated virus titers. The shown values are corrected for batch effects (See methods and **Table S4**). c, Bar-plot for cleaved RNA products and 28S rRNA measured with the BioAnalyzer system for the RNA samples from Calu-3 cells, before and after the viral infection for 24 h, with and without CaNDY treatments. Error bars, ±SD; *, p < 0.05; by Student’s t-test. d, RT-PCR results for *IFNB1* mRNA. e, Bar-plot for quantified IFNB1/ACTB in the infected cells based on RT-PCR results. Error bars, ±SD; **, p < 0.01; by Student’s t-test. f, A summary of the current study. **Figure 3-source data 1 (Separated File)** Original BioAnalyzer system result file for Figure 3C. **Figure 3-source data 2 (Separated File)** Original gel electrophoresis files for Figure 3D.

These results indicate that CaNDY treatment enhances the infection-dependent RNase L pathway, and the type I interferon pathway induced by the degraded RNAs, via a switching of *OAS1* splicing isoforms from *p42* to *p44a* in Calu-3 cells (Fig. 3f).

## Discussion

In this study, we show that the difference in the balance of *OAS1* splicing isoforms affects the infectivity of SARS-CoV-2 in cells via modulation of the innate immune response. This result strongly suggests that the *OAS1* intervening sequence (IVS) 5-1 G> A SNP that alters the *OAS1* splicing pattern is one of the important therapeutic targets for COVID-19. We recently reported on CaNDY as a potential therapeutic drug for cystic fibrosis caused by a point mutation in the *CFTR* gene (Shibata et al., 2020). In this study, we found that CaNDY treatment increased the yield of the *p44a* splice form (Fig. 2d, 2e, 2f) and made the Calu-3 cells more resistant to viral infection (Fig. 3b), suggesting that CaNDY treatment could overcome A-allele-derived vulnerabilities to SARS-CoV-2 infections.

The G-allele of SNP rs10774671 produces the *p46* splice variant, which possesses 56 amino acids at the C-terminus, compared to the 18 completely different amino acids present in the *p42* splice variant produced by the A-allele. This changes the localization of the OAS1 protein and its interacting partners (Kjær et al., 2014), which may be linked to the differential 2’-5’-oligoadenylate synthesis activities, RNase L activation, and interferon pathway activation. Several clinical trials that investigate the efficacy of an intervention of the interferon β pathway to affect the progression of COVID-19 have been conducted successfully (Hung et al., 2020; Monk et al., 2021; Shalhoub, 2020). On the other hand, inhibition of the interferon pathway is considered important to prevent cytokine storms in patients with severe COVID-19 (Nile et al., 2020). Since the *OAS1* IVS5-1 SNP can alter the response intensity of the interferon β pathway, stratification of COVID-19 patients depending on this SNP may assist in choosing the best course of interferon treatment.

It is reported that protective SNPs at *OAS1/OAS3*/*OAS2* loci originated in the Neanderthals (Zeberg and Pääbo, 2021; Zhou et al., 2021). Interestingly, the *OAS1* IVS5-1 A-allele, which was confirmed to be associated with the aggravation of COVID-19 in this study, was not found in the ancient human genome, the sequenced Neanderthal genome, or the Denisovan genome (Mendez et al., 2013). This suggests that the G to A mutation at the *OAS1* IVS5-1 position occurred at a relatively modern age in human history. The *OAS1* IVS5-1 A-allele has expanded to become a major allele in the population, especially in the Asian region (Fig. S4) (Marcus and Novembre, 2017). This suggests the possibility that this variant has some positive effects on human survival, trading off the effect of weakening resistance against viruses.

The results obtained in this study indicate that the *OAS1* IVS5-1 SNP is involved in the exacerbation of COVID-19 by eliciting changes in the *OAS1* splicing isoform balance. Individuals with the A-allele may have a higher risk for SARS-CoV-2 infection and its aggravation than those with the G-allele. However, we successfully demonstrate a way to overcome these genetic risks by modulating the splicing phenomenon. Splicing modulation decreased the virus infection rate 0.56 times in cell-based assays (Fig. 3b). This theoretically implies that the peak number of SARS-CoV-2-infected individuals will be reduced by 42%, if we can alter the *OAS1* splicing patterns for all the individuals carrying the *OAS1* IVS5-1 A-allele and boost the immunity of the entire population (simulation using the <u>S</u>usceptible, Infected and <u>R</u>emoved (SIR) model, see Methods for details). We hope that this approach based on genome-based precision medicine with a splicing modulator will contribute to the management of the COVID-19 pandemic.

## Acknowledgments

We would like to thank Editage [http://www.editage.com] for editing and reviewing this manuscript for English language.

## Author Contributions

K.I. conducted bioinformatics analysis. M.A., M.D., and R. K. conducted biological experiments relating cultured cells. Y.M., T.T., and T.N. conducted viral experiments. K.I., M.A., and M.H. wrote the paper.

## Supplementary Information

**Fig. S1.**
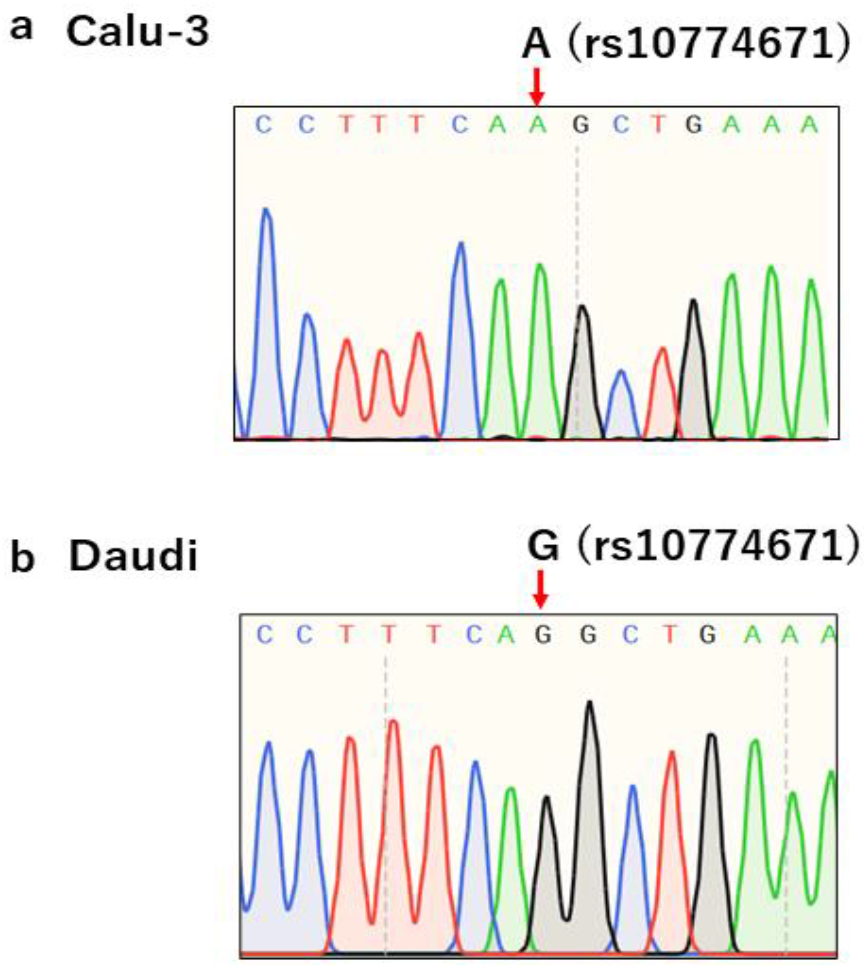
Genotyping of rs10774671 SNP. Results of the SNP genotyping for Calu-3 cells (a) and Daudi cells (b).

**Fig. S2.**
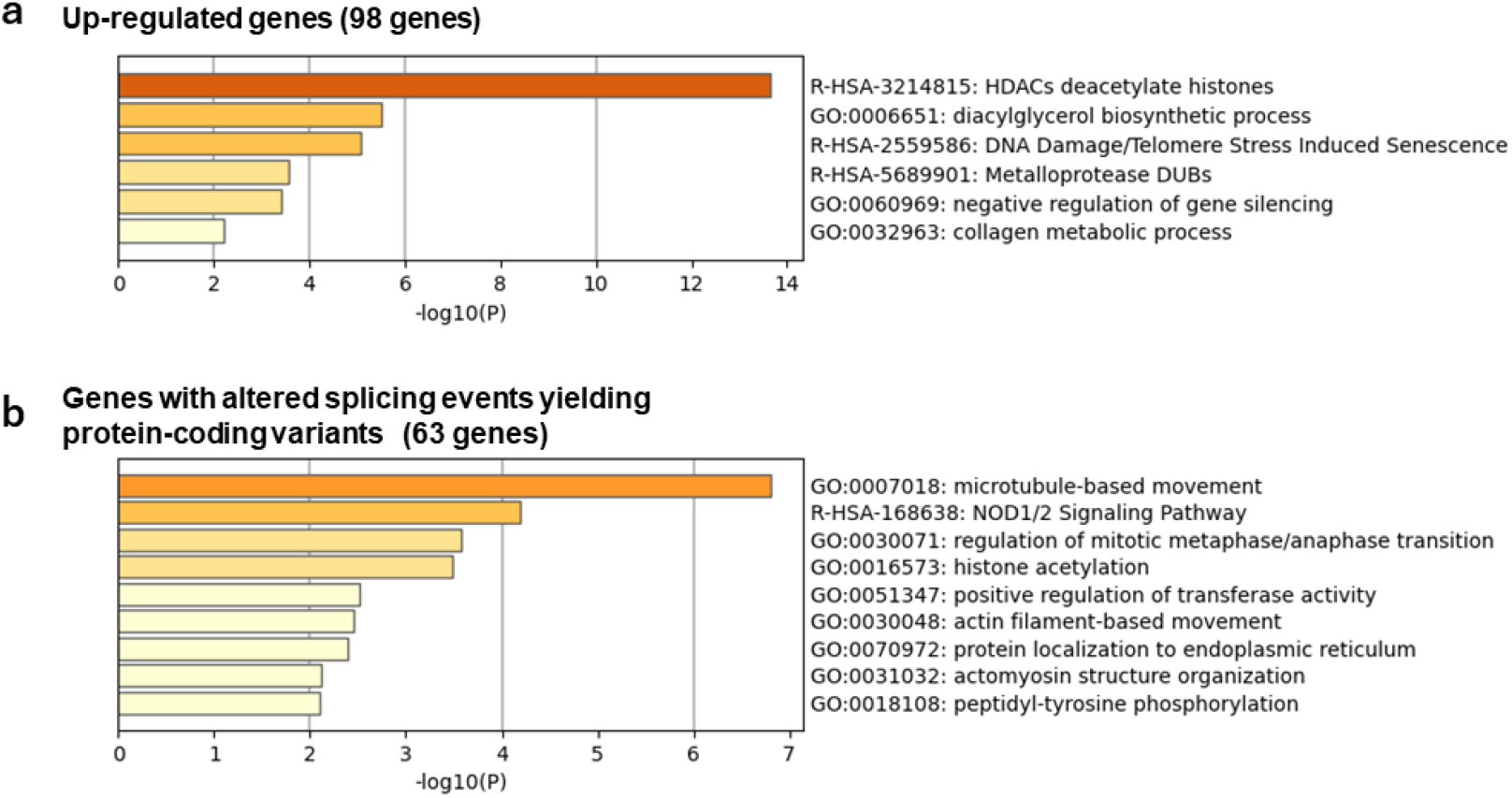
Pathway analysis for CaNDY-affected genes. Pathway analyses for 98 differentially up-regulated genes (a) and 63 differentially alternative spliced genes (b) in CaNDY-treated Calu-3 cells are shown. For the differentially alternative spliced genes, we used the splicing events that promote productive forms (see Methods for details). These analyses were performed with the Metascape web server (Tripathi et al., 2015)

**Fig. S3.**
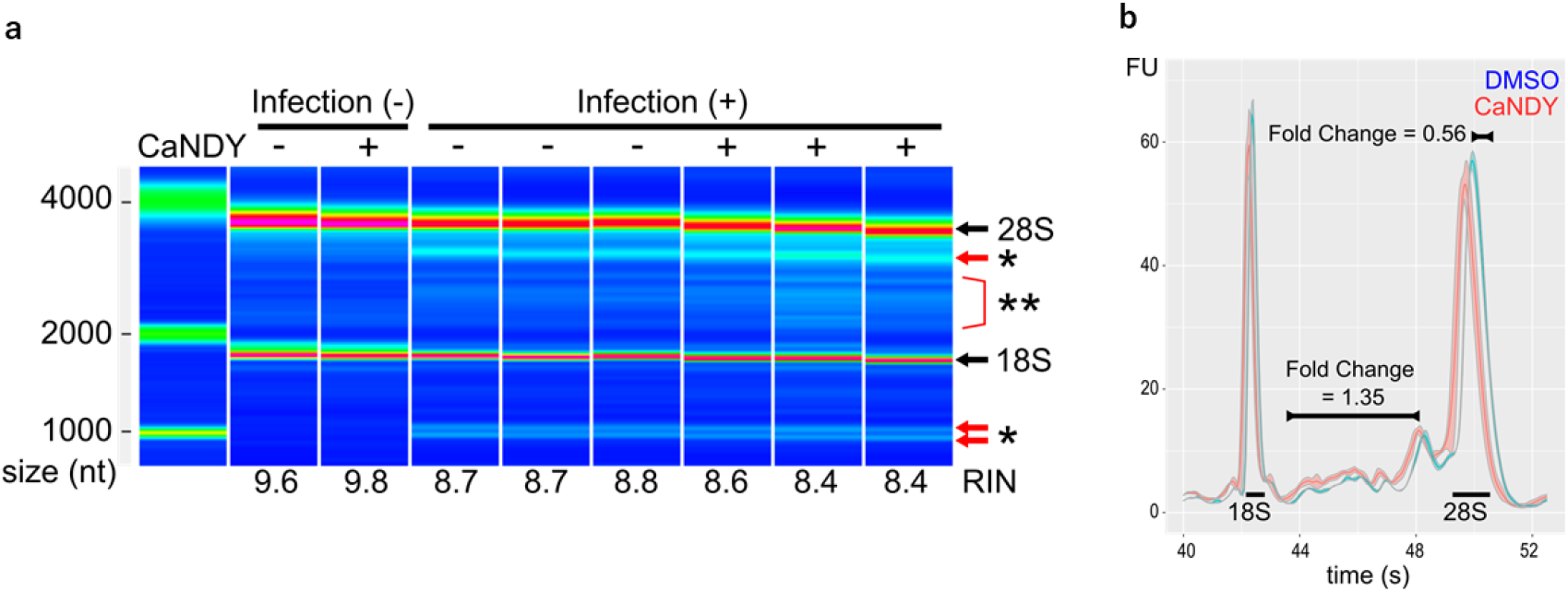
RNA degradation in SARS-CoV-2-infected cells. (a) A gel image was obtained based on the results from the Bioanalyzer. (b) Line graphs for fluorescence units according to migration times. Solid lines show the mean values and ribbons show the range of standard errors.

**Fig. S4.**
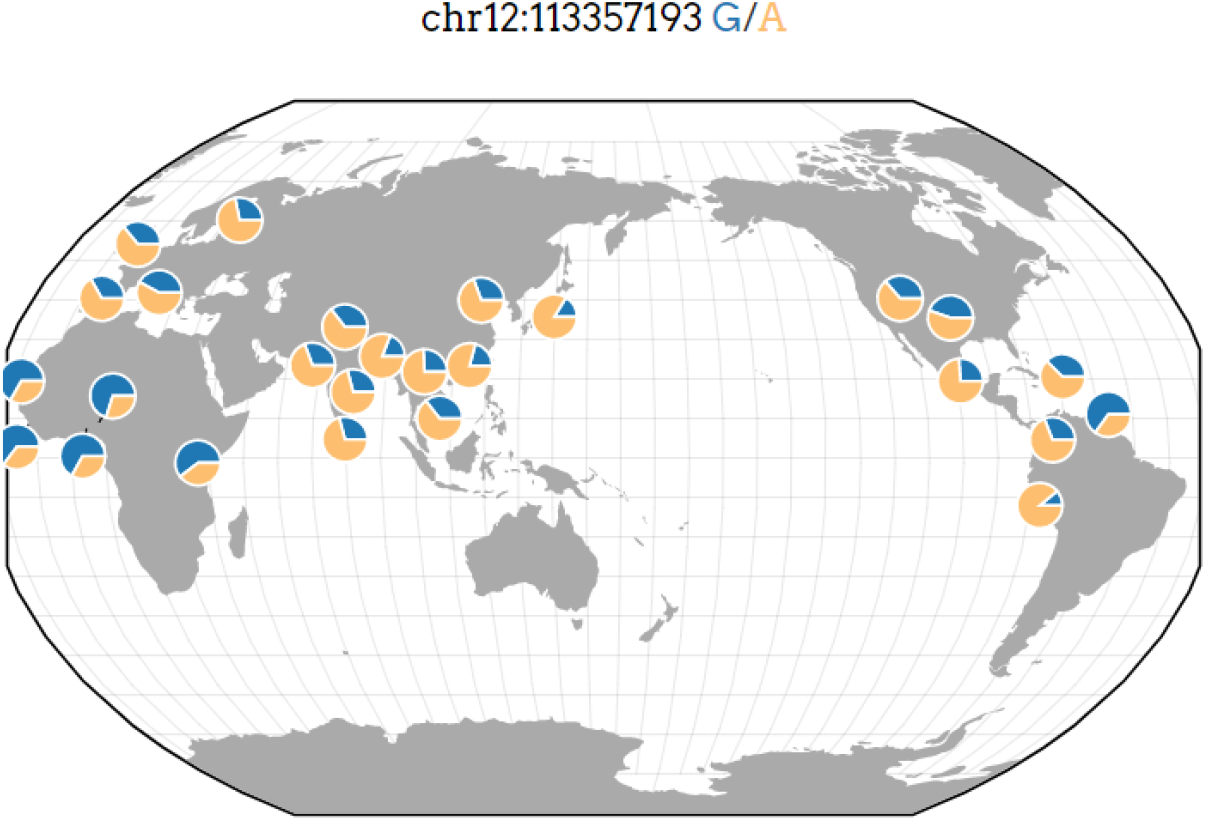
Allele frequency of the rs10774671 SNP. A world map showing the relationship between the location and allele frequency of the SNP. The position in the hg19 genome is shown. The map was obtained from the Geography of Genetic Variants (GGV) browser (Marcus and Novembre, 2017).

**Table S1.**
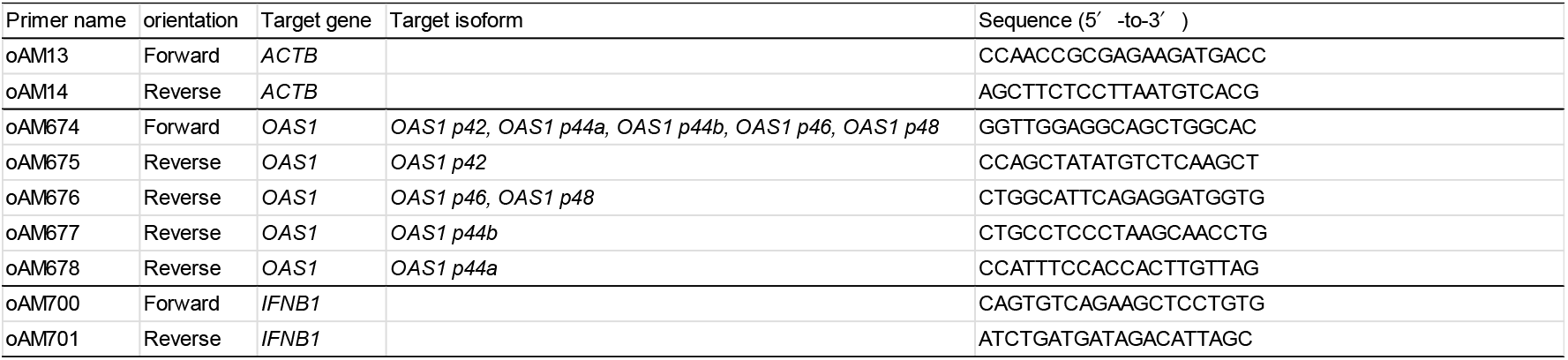
List of primers used in this study

Table S2. Up-regulated Genes in CaNDY-treated Calu-3 cells (Separate File)

Table S3. Altered splicing events yielding protein-coding variants in CaNDY-treated Calu-3 cells (Separate File)

**Table S4.** Raw and Batch-effect corrected results of the virus titer assay

| | Treatment | No. | $\log_{10}(\text{TCID}_{50}/\text{ml})$ | Batch-effect corrected |
| --- | --- | --- | --- | --- |
| **1<sup>st</sup> batch** | **CaNDY 0 $\mu$ M** | 1 | 7.75 | - |
|  |  | 2 | 7.67 | - |
|  |  | 3 | 7.75 | - |
|  | **CaNDY 10 $\mu$ M** | 1 | 7.50 | - |
|  |  | 2 | 7.50 | - |
|  |  | 3 | 7.25 | - |
| **2<sup>nd</sup> batch** | **CaNDY 0 $\mu$ M** | 1 | 6.75 | 7.60 |
|  |  | 2 | 7.25 | 7.92 |
|  |  | 3 | 6.50 | 7.44 |
|  | **CaNDY 10 $\mu$ M** | 1 | 6.75 | 7.60 |
|  |  | 2 | 6.50 | 7.44 |
|  |  | 3 | 6.45 | 7.41 |

